# Network Occlusion Sensitivity Analysis Identifies Regional Contributions to Brain Age Prediction

**DOI:** 10.1101/2022.10.31.514506

**Authors:** Lingfei He, Cheng Chen, Yaping Wang, Qingcheng Fan, Congying Chu, Junhai Xu, Lingzhong Fan

**Affiliations:** School of Computer Science and Technology, Tianjin Key Laboratory of Cognitive Computing and Application, Tianjin University, Tianjin 300350, China; Brainnetome Center, Institute of Automation, Chinese Academy of Sciences, Beijing 100190, China; National Laboratory of Pattern Recognition, Institute of Automation, Chinese Academy of Sciences, Beijing 100190, China; Chinese Institute for Brain Research, Beijing, China; CAS Center for Excellence in Brain Science and Intelligence Technology, Institute of Automation, Chinese Academy of Sciences, Beijing 100190, China; Sino-Danish Center, Beijing 100190, China; University of Chinese Academy of Sciences, Beijing 100190, China; University of Health and Rehabilitation Sciences, Qingdao, 266000, China

**Keywords:** Brain age, Convolutional neural network, Interpretability, Brain atlas, MRI

## Abstract

Deep learning frameworks utilizing convolutional neural networks (CNNs) have frequently been used for brain age prediction and have achieved outstanding performance. Nevertheless, deep learning remains a black box as it is hard to interpret which brain parts contribute significantly to the predictions. To tackle this challenge, we first trained a lightweight, fully CNN model for brain age estimation on a large sample data set (*N* = 3054, age range = [8,80 years]) and tested it on an independent data set (*N* = 555, mean absolute error (MAE) = 4.45 years, *r* = 0.96). We then developed an interpretable scheme combining network occlusion sensitivity analysis (NOSA) with a fine-grained human brain atlas to uncover the learned invariance of the model. Our findings show that the dorsolateral, dorsomedial frontal cortex, anterior cingulate cortex, and thalamus had the highest contributions to age prediction across the lifespan. More interestingly, we observed that different regions showed divergent patterns in their predictions for specific age groups and that the bilateral hemispheres contributed differently to the predictions. Regions in the frontal lobe were essential predictors in both the developmental and aging stages with the thalamus remaining relatively stable and saliently correlated with other regional changes throughout the lifespan. The lateral and medial temporal brain regions gradually became involved during the aging phase. At the network level, the frontoparietal and the default mode networks show an inverted U-shape contribution from the developmental to the aging stages. The framework could identify regional contributions to the brain age prediction model, which could help increase the model interpretability when serving as an aging biomarker.

## Introduction

Previous studies have shown that the human brain undergoes complex structural changes during development and aging, including widespread synaptic pruning and myelination from early life through puberty and neurodegenerative processes in later life [1]. In addition, research has shown that global gray matter decreases with age but that regional gray matter exhibits heterogeneous age effects [2, 3]. For example, the gray matter volume in the frontal and parietal lobes and some temporal lobe regions substantially decreased with age [4-7]. In contrast, some subcortical structures, such as the caudate and hippocampus, showed nonlinear developing patterns [8, 9]. As age increases, the influence of the aging process on individuals is also heterogeneous due to genetic and environmental effects [10, 11]. Potentially, the extent to which someone deviates from healthy brain aging trajectories could indicate underlying problems in outwardly healthy people and relate to the risk of cognitive aging or age-associated brain disease. Hence, reliable biomarkers of brain aging could be of great neuroscientific and clinical value [12].

Recent research has extensively studied the possibility of predicting brain aging from structural magnetic resonance imaging (MRI) brain scans [12-14]. Brain age has been recognized as a reliable brain development and aging biomarker [22]. By predicting the difference between individual brain age and chronological age, we can infer the risk of neurodegenerative diseases and even mortality [15]. Usually, brain age prediction is conducted through traditional machine learning methods requiring manual anatomical feature extraction [16-20] to fit a model with specific hypotheses. Although the increase in the number of available neuroimaging datasets and the improvements in computing resources have accelerated the development of this field, feature extraction is still time-consuming and knowledge-dependent, and the interpretation of the model can only rely on the extracted local features, which might lose potential spatial and global context information [21]. As such, using deep learning to predict brain age is now taking the stage [22]. Deep learning frameworks overcome the shortcomings of machine learning in that they can directly leverage 3D images without losing spatial feature information and obtain relatively higher precision results. For example, Cole et al. utilized a 3D convolutional neural network with a fully connected layer to predict brain age from a large dataset of healthy subjects aged 18–90 years and found that convolutional neural networks (CNNs) had an excellent age prediction performance with a mean absolute error (MAE) = 4.16 [22]. Recently, Jonsson et al. combined a 3D CNN with a residual neural network to build a brain age prediction model [23]. Because the residual connections improved the network structure and the dataset size increased to tens of thousands of cases, the model gained remarkably enhanced prediction accuracy and generalization performance.

Although deep learning provides highly accurate age prediction, the size and complexity of the network still make it a black box, and it is difficult to identify the features that have an essential impact on the prediction. Several recent studies have attempted to interpret or visualize the intermediate representations of CNNs. For example, a pioneering inverse-structural method, the CNN visualization proposed by Matthew et al. [24], used deconvolution and unpooling to transform the hidden representation of a network model into an image with practical and interpretable meaning, realizing feature visualization and helping us understand what each layer of the CNN had learned. After that, some gradient-based methods took the lead in interpretative visualization. Representative gradient-based methods, such as gradient-weighted class activation mapping (Grad-CAM) [25], are commonly used to interpret the classifications produced by CNNs and have been previously used in CNN-based medical image analysis [26] to marry potential disease pathology with classification findings to generate a series of age-averaged activation maps to study the age-specific patterns of the underlying basis for age estimation [27]. Compared with a previous CAM [28], Grad-CAM can visualize the CNN of any structure without modifying the network structure or retraining and can distinguish the image position without attention. In addition, an outstanding gradient-based method called SmoothGrad [29] can create a population-based explanatory graph to explain the CNN model [30]. In this method, a given input image is first distorted using random noise sampled from a normal distribution N(μ = 0,σ = 0.1). Then, the partial derivative of each voxel is computed for the trained model’s output. This method has been demonstrated to be able to capture the CNN training process [31]. Furthermore, the recent work by He et al., in which they successfully implemented a two-channel brain age prediction model based on global and local information by utilizing the self-attention mechanism [32] in the Transform model [33], is notable in that the prediction accuracy reached the optimal level. Because the model had learned local information, it could generate the corresponding probability heat map for the explanation.

However, these explanatory methods still have problems on several levels. For example, some background units may be regarded as significant components of the effects on classification, potentially adding confounding variables. As a result, the degree of contribution of specific brain regions, which is the focus of the explanation, can neither be accurately located nor be targeted so that it can be quantified. Furthermore, because the results are often vague or with a priori bias, the methods that have been shown to be effective on naturalistic images may not retain good interpretability on actual neuroimages, so their results may not be persuasive enough to be generalizable [34-38].

Given the above considerations, in this study we aggregated a large, heterogeneous dataset of structural neuroimaging from multiple publicly available data sources to explore the change patterns of age prediction as they relate to multiple brain regions throughout the human lifespan. We filtered the data to distribute the sample size evenly to prevent a biased model. Subsequently, we leveraged these data to implement a 3D fully convolutional neural network for brain age prediction. This process reduced the parameter size and increased the speed of our training process without losing accuracy. Finally, we employed network occlusion sensitivity analysis (NOSA) [24] from the field of computer vision and utilized regions from the human Brainnetome Atlas [39] to reveal the most relevant brain regions for age prediction. We also proposed a quantitative index, i.e., an importance score (IS), for each brain region to facilitate the observation and explanation process. Further, using independent data sets, we investigated the reproducibility of IS as a metric and observed the distribution patterns of this metric over age across brain regions.

## Materials and methods

### Datasets

Building an adequate neuroimaging dataset for age prediction over a wide range of ages is necessary to explore the patterns of brain development and aging across the lifespan. In addition, a uniformly distributed data set can train the model to distinguish the structural differences underlying the neural information to produce unbiased results instead of giving a specific value near the center of the data bias. Therefore, we filtered the following data to distribute the sample size uniformly.

In this work, we constructed a training set and an independent test set with an age range of 8-80 years old and with a uniform distribution. The training set contained a sample of 3054 T1-weighted MRI brain scans of healthy individuals (male/female = 1478/1576, mean age = 42.49±13.47, age range 8-80 years) from seven publicly accessible databases. The independent test set was derived from the Human Connectome Project (HCP) data set (*n* = 555). All scans were acquired for studies that had been approved by the local institutional review boards, research ethics committee, or human investigation committees. We only included participants without major neurodegenerative or psychiatric diseases. All the T1-weighted MRI data were acquired on either a 1.5 T or 3 T scanner using standard T1-weighted sequences. The detailed information and age distribution of all the data used in our work are shown in Table 1. Moreover, we constructed a multisite dataset (*n* = 313) derived from the Dallas Lifespan Brain Study (DLBS) (male/female = 101/70, mean age = 49.85±17.44, age range 20-80 years) and the Philadelphia Neurodevelopmental Cohort (PNC) (male/female=23/19, mean age=13.14±3.14, age range 8-20 years; not included in the training set) databases by the same sampling strategy as an additional test dataset for verification.

**Table 1.** Dataset information and age distribution of the training set and the HCP independent test set.

| Dataset | *n* | Age range | Age mean (±SD) | Gender (M; F) |
| --- | --- | --- | --- | --- |
| IXI | 540 | 20-80 | 48.10 (±16.16) | 234; 306 |
| SALD | 494 | 19-80 | 45.18 (±17.43) | 307; 187 |
| NKI | 422 | 18-80 | 46.67 (±18.56) | 141; 281 |
| CoRR | 370 | 18-79 | 40.69 (±20.08) | 196; 174 |
| UKB | 697 | 45-80 | 61.59 (±11.51) | 351; 346 |
| PNC | 398 | 8-20 | 13.54 (±3.43) | 175; 223 |
| 973 | 133 | 30-40 | 34.30 (±3.17) | 74; 59 |
| HCP (Test set) | 555 | 8-80 | 44.18 (±20.48) | 293; 262 |
| Total (*N*) | 3609 | 8-80 | 41.78 (±13.85) | 1771; 1838 |

### Data Preprocessing

All T1w MRI data were processed using the CAT12 toolbox (http://www.neuro.uni-jena.de/cat/) in the Statistical Parametric Mapping toolbox (SPM12, https://www.fil.ion.ucl.ac.uk/spm/) in MATLAB R2017b (University College London, London, UK). Based on their tissue classification, the T1-weighed MR images were segmented into gray matter, white matter, and cerebrospinal fluid, but only gray matter was used in this study because of the high correlation between gray matter and the brain aging process [40-42] and to enable methodological comparability with previous studies [43, 44]. Then, the gray matter maps were spatially normalized into the Montreal Neurosciences Institute (MNI) space using the DARTEL algorithm. After preprocessing, the gray matter images had dimensions of 121 × 145 × 121. Finally, a 4 mm full-width half-maximum Gaussian filter kernel was used to smooth the modulated gray matter images. These smoothed gray matter images were used in the subsequent analysis.

In Table 1, the statistical data is presented using the name of the fMRI dataset (in the Dataset column), the number of subjects (*n*), age range, age mean with standard deviation, and gender. The settings for acquiring these datasets were different, so the resulting dataset was heterogeneous. After filtering using the specific strategy described in the text, the training and test datasets were uniformly distributed by their age ranges and thus had the lowest possible bias.

### Convolutional neural network

Our 3D CNN architecture referenced a recent excellent brain age prediction model [45], which was based on visual geometry group classification architecture [46] and a fully convolutional network [47]. The input for our model was a 3D gray matter image with dimensions of 121 × 145 × 121, and the output contains 73 digits representing the predicted probability that the subject’s age falls into a one-year age interval between 8 to 80. Specifically, the architecture of the CNN model included five repeated blocks, and the structure of each block included 3 × 3 × 3 convolutional layers (with a stride of 1 and padding of 1), followed by a 3D batch-normalization layer, a rectified linear unit (ReLU) activation function and a 2 × 2 × 2 max-pooling layer (with a stride of 2). Next, there was a 1 × 1 × 1 convolutional layer (followed by a 3D batch-normalization layer and a rectified linear unit (ReLU) activation function), which can further increase the nonlinearity of the model without changing the output size of the feature map. Finally, we used average pooling, a 1 × 1 × 1 convolutional layer, a flattening operation, and a SoftMax layer to replace the fully connected layers to generate the probability distribution of the predicted age. The channel numbers used in each convolution layer were [32, 64, 128, 256, 256, 128, 73], and the weighted sum of each age label was calculated to make the final prediction:

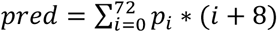

where *p*_*i*_ stands for the probability predicted for the *i*^th^ age class, (*i*+8) stands for *i*^th^ age class label. The details of the CNN architecture are shown in Figure 1(A).

**Figure 1.**
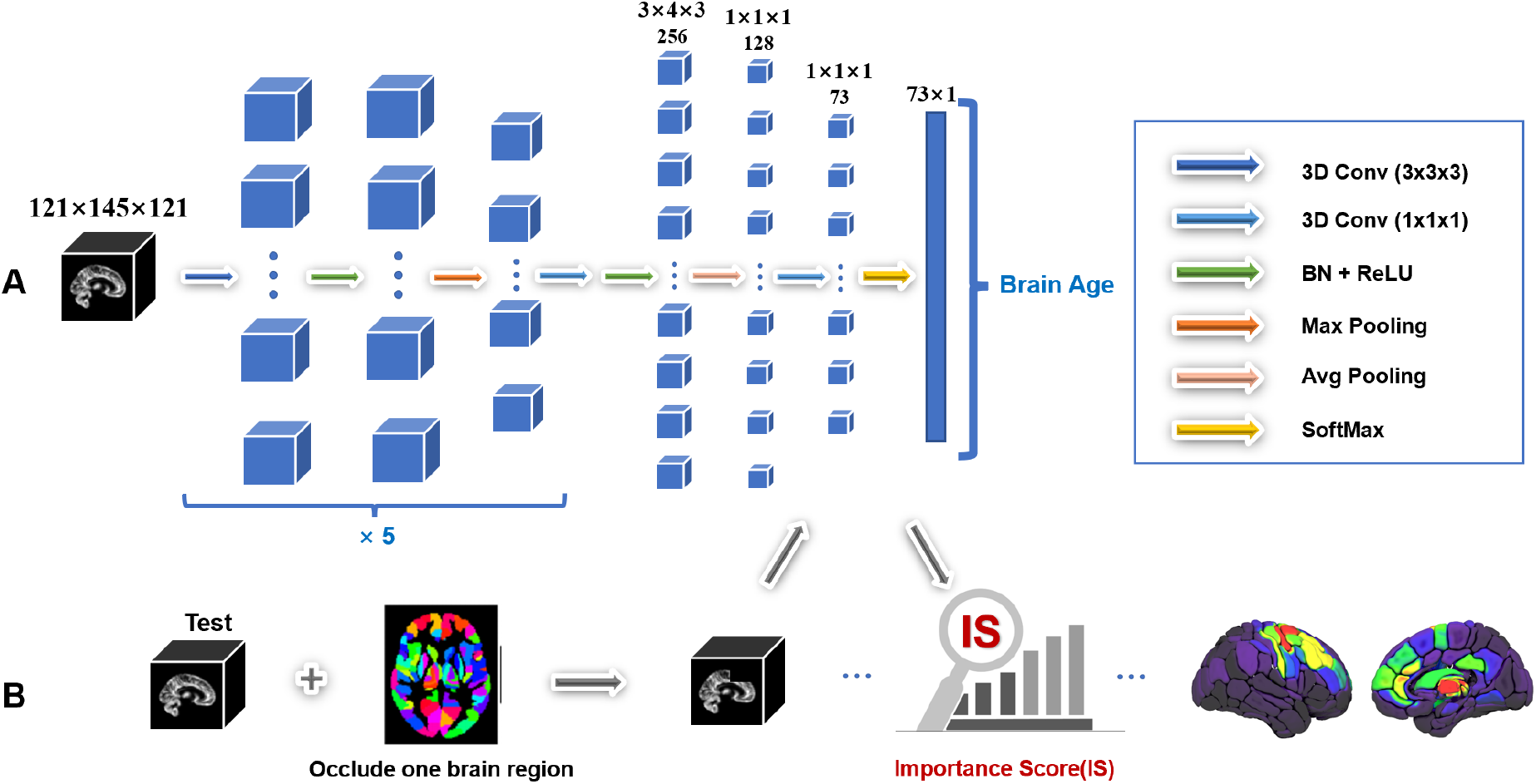
(A) The architecture of our 3D fully convolutional neural network. The black box represents the input structural MRI image, the blue boxes represent feature maps, and the first three layers are repeated five times. The arrows indicate 3D CNN operations, and the model outputs the probabilities of 73 age categories weighted by their corresponding ages and summed up to produce the final brain-predicted age. The texts above the 3D feature maps are the dimensional annotations— the first line denotes the feature dimensions while the second denotes the channel. All batch dimensions are hidden for ease of understanding. (B) The network occlusion sensitivity analysis (NOSA) method in our work. According to the human Brainnetome Atlas definition, one brain region was occluded as the test input in the testing phase. After obtaining the predicted ages of the initial input and test input inference by the above CNN model, an IS of the target brain region was calculated to further analyze the brain aging process.

### Model training and evaluation

We used Pytorch to implement the algorithm and train the model with ten-fold cross-validation. During the training process, the cross-entropy loss was used as the loss function, and the model was optimized using stochastic gradient descent (SGD) [48]. We set the mini-batch size to 12, the learning rate to 0.01 with a constant decay of 0.3 after every 50 epochs, the weight decay to 0.001, and the number of epochs to 300. Then, the initialization strategy [49] was used to initialize the weights. Finally, the model with the lowest average validation mean absolute error (MAE) between the real and predicted age was selected as the optimal model.

In the subsequent testing phase, we evaluated the generalization ability and effect of the model by calculating the MAE, the Pearson correlation coefficient (*r* value), and determination coefficient (*R*^2^) on the HCP independent test set and the multi-center test set.

### Analysis of the brain regions related to brain age prediction

Before we applied the human Brainnetome Atlas as the basis of the NOSA method, some comparative experiments were conducted to explore the influence of the atlas (See supporting text and Fig. S1 included in the supplementary materials). In the end, we used the human Brainnetome Atlas with the NOSA method to explain our model and determine the brain regions relevant to brain age prediction. Specifically, in the testing phase, we occluded one brain region at a time according to the 246 brain regions defined by the human Brainnetome Atlas. We recalculated the MAE and compared the MAE changes before and after:

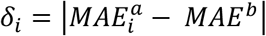

where *MAE*^*b*^ and 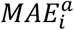 respectively represent the testing mean absolute error of our entire HCP independent test set before and after the *i*^th^ brain region is occluded. A higher *δ* value indicates that this brain area plays a more critical role in brain age prediction. Subsequently, we normalized the 246 *δs* and defined them as the importance scores (ISs) of the 246 brain regions:

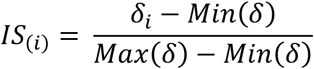

where *IS*_(*i*)_represent the importance score for the *i*^th^ brain region, and *Max*(*δ*) and *Min*(*δ*) represent the maximum and minimum of the 246 *δs*, respectively. A NOSA test is shown in Figure 1(B).

To verify the results obtained on the HCP independent test set, we repeated the work on the uniformly distributed multi-center dataset. In addition, we reproduced the GBP method [50] as a methodological comparison. The GBP method is generally regarded as a standard visualization method for exploring the interpretability of a CNN in that this method determines the relevance of each pixel for the classification decision by calculating the gradient of the class output score for the input image. Here, we calculated the gradient of each voxel backpropagation using the GBP method on the trained model to obtain the salience map and then calculated the average gradients of the voxels belonging to every brain area in the human Brainnetome Atlas. The above process was repeated ten times, and finally, each average value of the gradient was taken as the contribution of the corresponding brain area.

### The pattern of changes in the IS of brain regions with age

To observe changes in the importance of brain regions related to brain age prediction, we divided the HCP independent test into seven age groups. The age spans of each group were: [8,20), [20,30), [30,40), [40,50), [50,60), [60,70), [70,80], and the sample size distribution of each group was uniform, as shown in Table 1. Then we calculated the IS of the 246 brain regions in each group and overlayed it on the human Brainnetome Atlas standard space template for a visual analysis. We also performed a region-wise Spearman correlation and polynomial regression analysis and identified the most relevant regions that also had a high IS across the lifespan (See Supplementary Material, Fig. S2-4).

### IS patterns at the brain network level

We calculated the IS values of 246 brain regions of all the individual subjects (*n* = 555) in the HCP independent test dataset for a network analysis. This generated an IS matrix with a dimension of 555 × 246, which means that each brain region had a coverage IS change curve for 8-80 years old. Then, we used the 7-network resting state network parcellation defined by Yeo and colleagues based on data from 1000 subjects [51] and the IS matrix obtained above to investigate the changes in the contribution of each brain region at the network level. Specifically, based on the 7-network divisions, we obtained the average value of the ISs of all brain regions belonging to the same network and used it as the IS value of the network. This generated an IS matrix with a dimension of 555 × 7 at the network level. After that, to further observe the IS changing pattern of each network, we also applied polynomial regression to quantify the relationship between IS and chronological age for each network separately based on ordinary least squares (OLS) calculations.

### The lateralization of the IS of brain regions

It is well known that the left and right hemispheres have specific functional divisions. There is the phenomenon of lateralization, in which the left hemisphere is mainly responsible for processing abstract information, such as text and data, whereas the right hemisphere is primarily responsible for processing specific information, such as sound and images [52]. Since the model above is based on whole-brain image training, one hemisphere may be affected by the other hemisphere when studying lateralization. To explore the lateralization of the importance of brain regions, we used the same data set and brain age prediction model framework with the same configuration as above to train two additional brain age prediction models based on the left and right hemispheres.

We changed the input whole-brain images into hemispheric images to train the brain age prediction model on each hemisphere. We first made separate masks of the left and right hemispheres based on the partitions of the Brainnetome Atlas. The value of the reserved hemisphere was one, and the value of the other hemisphere was 0. Since all images were registered to standard space, the whole brain image of a subject could be multiplied by the corresponding hemispheric mask to obtain the hemispheric image of the subject. Second, since the value of one hemisphere was set to 0, to speed up the model’s training we cut out the deleted hemisphere so that the dimension of the input image changed from 121 × 145 × 121 to 61 × 146 × 121. In addition, it should be noted that since our model architecture is based on a fully convolutional neural network, changing the input dimension of the model did not affect the previous overall architecture.

## Results

### Brain age prediction

We used ten-fold cross-validation to evaluate our CNN model and selected the model with the lowest average validation MAE as our final optimal model. On the validation set, the optimal CNN model achieved a MAE = 2.85, Pearson correlation coefficient *r* = 0.98, and determination coefficient *R*^2^ = .96. Moreover, the test performance achieved MAE = 4.45 years, *r* = 0.96, *R*^2^ = .92 on the HCP independent test set and MAE = 4.40 years, *r* = 0.96 and *R*^2^ = .92 on the multi-center test set (see Fig. 2A).

**Figure 2.**
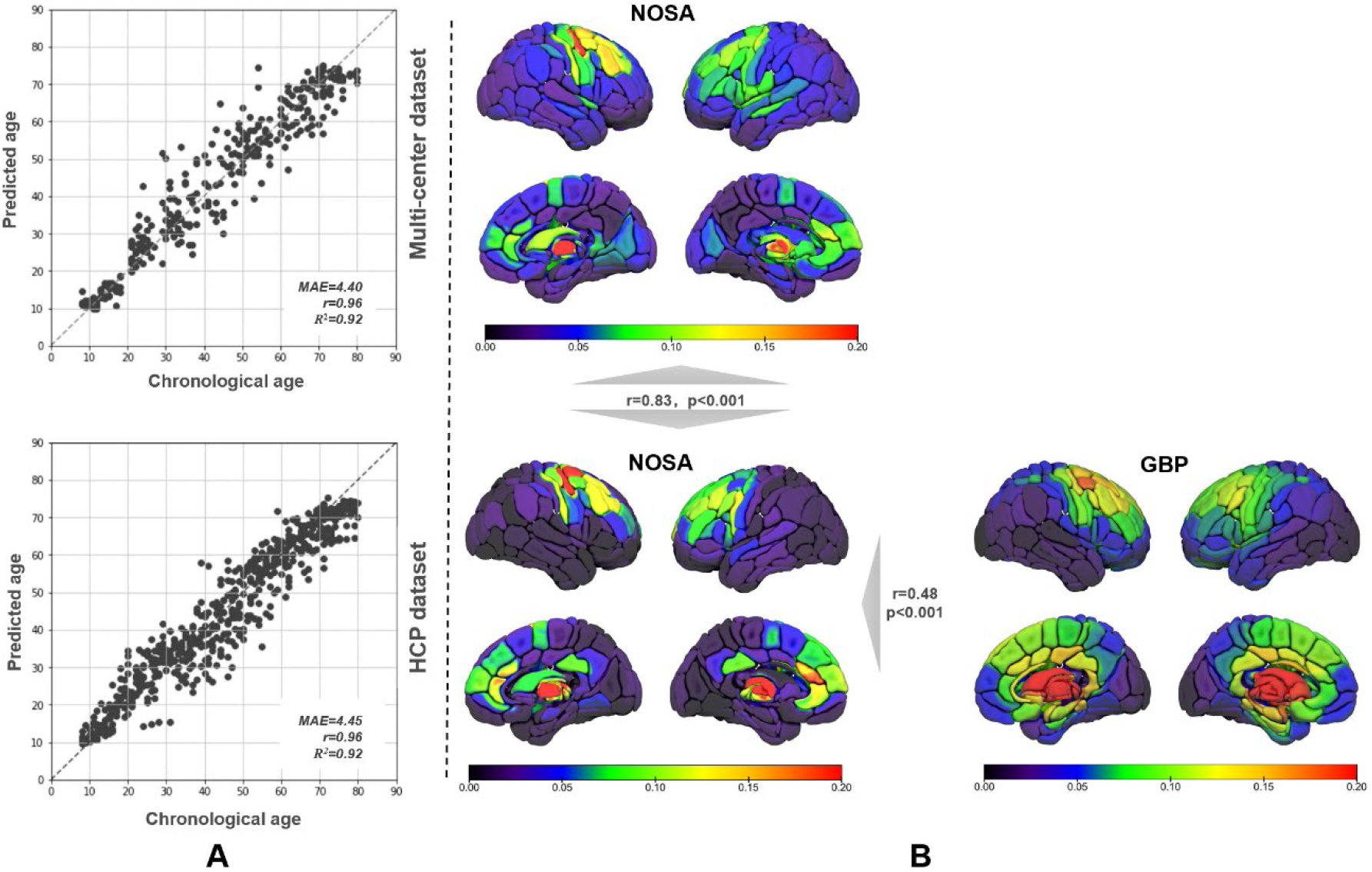
Test performance of our model and stability verification and methodological comparison between GBP and NOSA. (A) Estimated age versus chronological age with predictive metrics. Our model showed a good and stable performance on both test datasets. (B) The distribution is plotted on the standard brain template, including IS calculated by our NOSA method on both test datasets and salient values calculated by the GBP method on the HCP dataset. The NOSA method showed a significant correlation with the GBP method on the HCP dataset, and its stability was verified on the multi-center dataset.

### Brain regions that play an essential role in predicting brain age

In the entire HCP independent test set, we used the proposed network occlusion sensitivity analysis method to obtain the IS values of 246 brain regions based on the definitions of the human Brainnetome Atlas. Then we sorted the 246 IS values to observe which brain regions would be of greater importance. The top 15 IS values and brain regions information are shown in Table 2. Throughout the human lifespan (8-80 years in our work), we found that the dorsolateral, dorsomedial frontal cortex, anterior cingulate cortex, and thalamus made the greatest contributions to age prediction. Of these, the score of the thalamus region was generally very high, which makes it seem particularly important. This result is consistent with the findings of recent studies [21, 26].

**Table 2.**
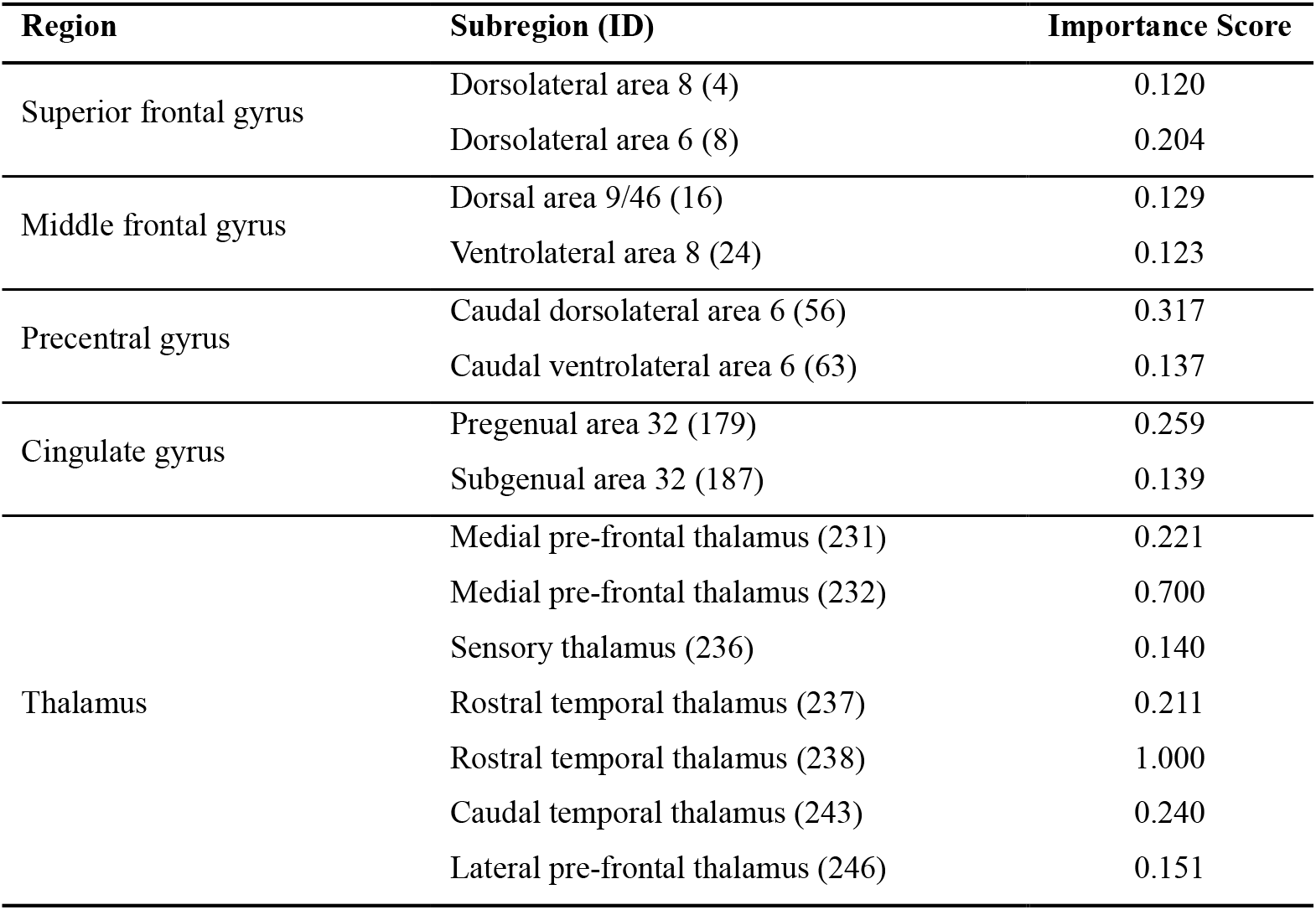
The top 15 IS values and brain regions information.

The results are presented as the name of each brain region, the name of the corresponding subregion with ID, and the IS of each subregion. The human Brainnetome Atlas defines all regions and subregions. The IS values were obtained by repeated measuring and rounded to three decimal places. We noticed that the score of the thalamus region was strikingly high, making it seem particularly important.

### Comparing the results from using the multi-center test set and the GBP method

The relationship between the estimated age and chronological age is shown in Figure 2 (A, B). The Pearson correlation coefficient of the results obtained by the NOSA method in the HCP and the multi-center data set was 0.83, *p* < .001. The results produced by the two test sets tended to be consistent, which increases the reliability of the results. In another set of comparative experiments, the Pearson correlation coefficient of the NOSA method and the GBP method on the HCP test set was 0.43, *p* < .001. For the GBP method, we found that the brain regions that had an important influence on the prediction of brain age were mainly the frontal cortex and thalamus, similar to the results obtained by the NOSA method, Figure 2 (B).

### The pattern of changes in the importance of brain regions with age

To explore how the importance of brain regions changed with age, we divided the HCP independent test set into seven groups with increasing ages, and each group had the same age span except for the first group—[8, 20), [20, 30), [30, 40), [40, 50), [50, 60), [60, 70), [70, 80]. We first calculated the MAE of each group and observed its changes. The MAE values for each age group were 2.483, 5.220, 4.408, 5.833, 4.484, 3.461, and 4.324, respectively. These values show that the model had a lower MAE for the young and old groups, while the MAE was higher in adulthood. This phenomenon is not difficult to understand, mainly because the changes in brain regions are particularly obvious in the young and aging stages, and the model is able to learn easily. In the adult stage, the changes were not noticeable, making the model unable to learn the most outstanding features. This result shows from a different perspective that our model could well capture the changes in brain regions.

Subsequently, we used network occlusion analysis for each age group on the optimal model and generated 246 IS values corresponding to the 7 age groups. The distribution of the IS values of the 7 groups on the human Brainnetome Atlas is shown in Figure 3. From the results of these seven groups, we found that different brain regions showed different patterns of brain age prediction at the different age levels. Overall, the brain regions in the frontal lobe became the essential predictors in both the developmental and aging stages. In contrast, the lateral and medial temporal brain regions mainly showed contributions in the aging phase. Most brain regions seemed to have no apparent changes in the mature stage, which aligns with general biological knowledge. In addition, more interestingly, we found that the thalamus made a significant contribution in all stages. Finally, we performed the same grouping experiment on the GBP method, but that method did not capture the patterns of brain regions that change with age. In contrast, the results between the groups were highly correlated with almost no differences.

**Figure 3.**
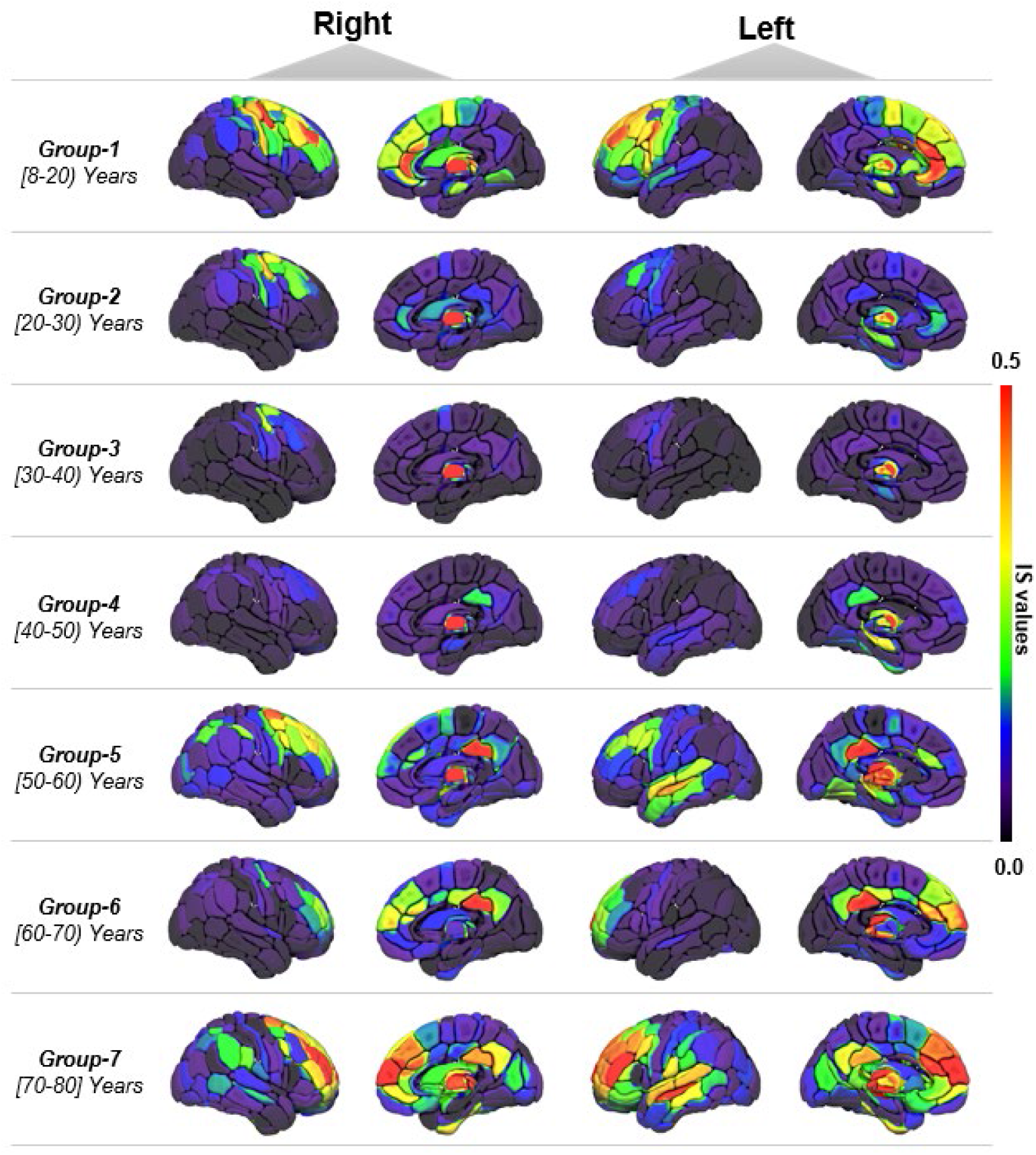
The distribution of IS values of the seven age groups on the standard brain template. The result shows divergent patterns in brain age prediction at varying age levels on both hemispheres. The closer the color is to red, the greater the importance of the brain area for brain age prediction. Overall, brain regions in the frontal lobe became the essential predictors in both the developmental and aging stages. In contrast, the lateral and medial temporal brain regions mainly made contributions in the aging phase.

After that, to obtain accurate insights, we also performed correlation and regression analyses (See supporting text and Fig. S2-4). We noticed that the regions in the thalamus that had salient ISs throughout the whole life also had the highest correlation with age and a broad correlation with other integrated parts (See Fig. 4), which reconfirmed the importance of the thalamus in the brain aging process.

**Figure 4.**
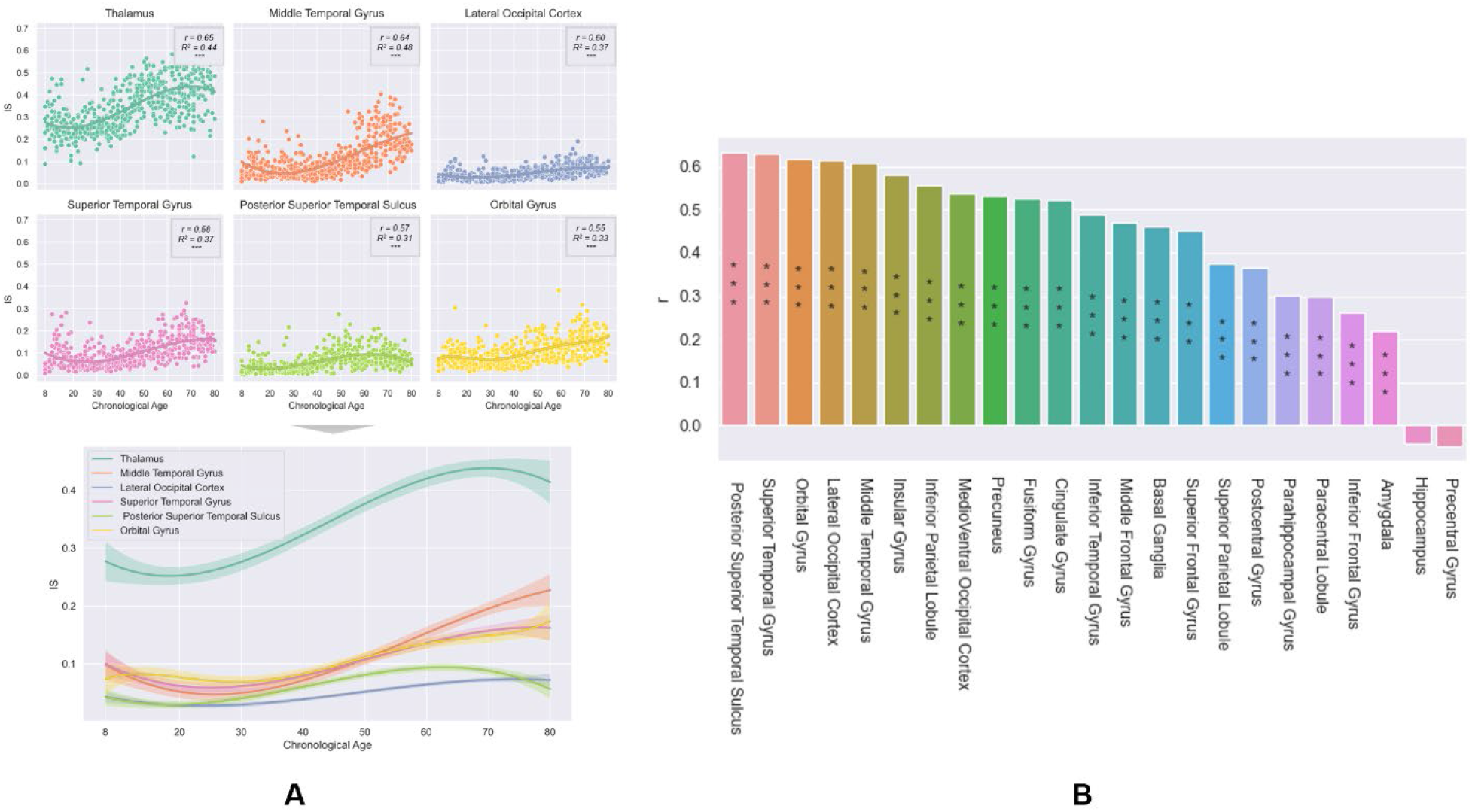
Regional correlations of the thalamus. (A) Integrated comparison with the other most age-related regions. The upper six subplots represent the most age-related integrated parts ordered by their correlation coefficients. The annotations on the upper left in each subplot are the correlation coefficients *r*, determination coefficient *R*^2^, and the significance level of *r*. The lower chart is the aggregated result. All regression functions passed the significance test with *p* < .01. (2) The ordered IS correlations between the thalamus and other regions. All the asterisks in both charts are significance levels (‘***’ means *p* < .01). We observed that the regions aggregated in the thalamus gained top correlations with age and high overall ISs. In addition, the result showed a high consistency in the changes between the thalamus and almost all the other integrated parts, especially the most age-related regions, revealing the critical role of the thalamus during aging.

### IS values at the 7-network level

After that, we calculated the IS values of the seven networks defined by Yeo on 555 subjects and observed their changes. As shown in Fig. 5, the fitted regression lines can broadly reflect the IS variation tendency of each network with a high degree of statistical significance. (*p* < .01). All networks had very low IS values in the maturity stage. At the same time, only the frontoparietal network (FPN) and the default mode network (DMN) significantly changed during the development and aging stages. These results indicate that, compared with other networks, these two networks play an essential role in predicting brain age during development and aging.

**Figure 5.**
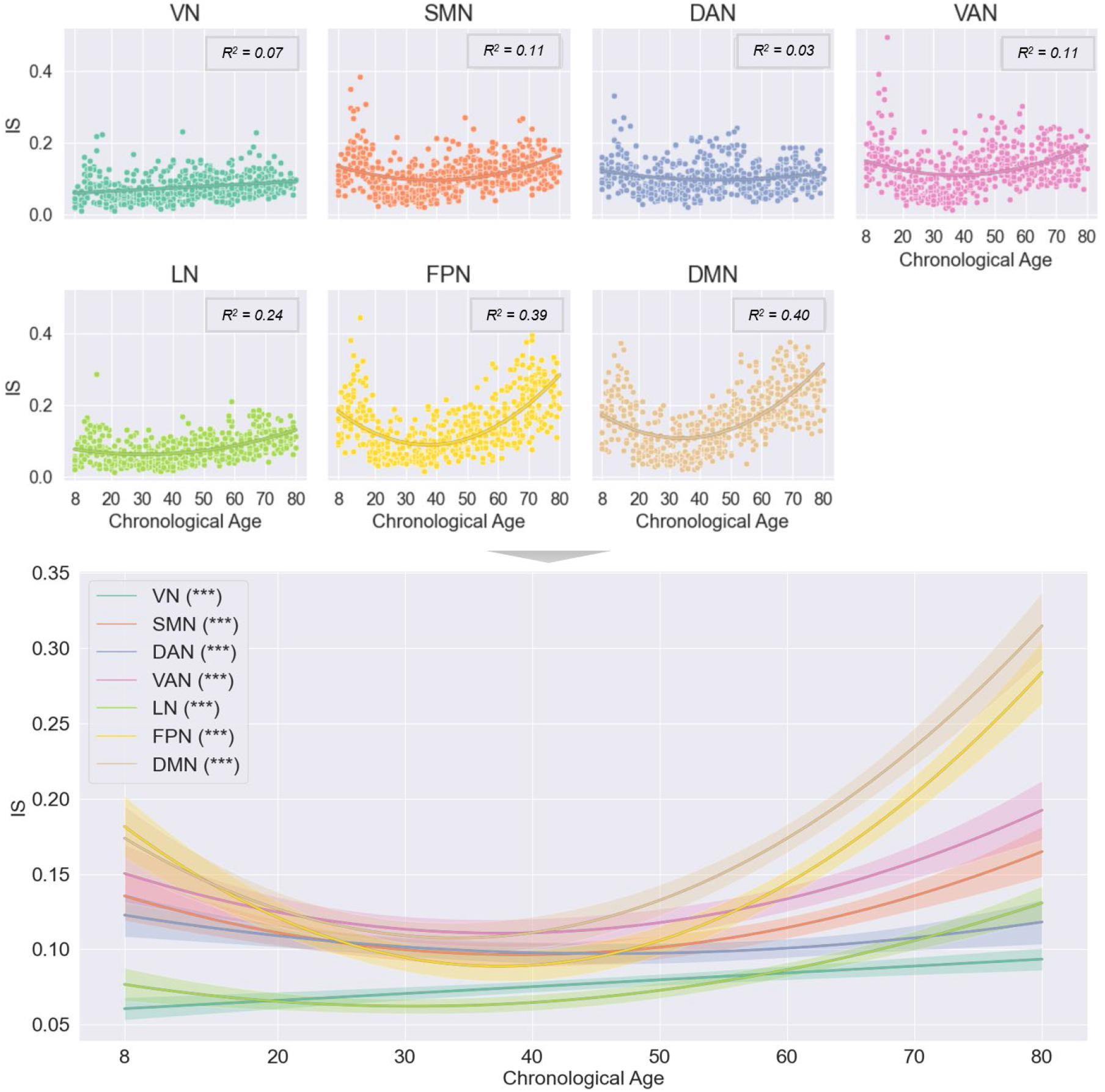
Changes in IS values of 7 networks with age. We categorized the brain regions into different functional brain networks and then observed various brain networks with varying patterns of changes in IS values with age. The subplots above denote the individual IS distributions of each network with increasing age. The optimal polynomial regression line in each subplot reveals the IS variation tendency for the corresponding network during aging. *R*^2^ denotes the determination coefficient. The aggregated line chart with error bands below reflects the overall comparison between the 7 networks, and the three asterisks behind each network annotation mean that every parameter in the corresponding regression function was at a high degree of statistical significance *p* < .01. This shows that the FPN and DMN changed significantly in the development and aging stages, playing an essential role in predicting brain age.

### Lateralization of the importance of brain regions

Figure 6 (A) shows the results of two optimal brain age prediction models based on the left and right hemispheres, including the MAE, *r*, and *R*^2^ values between the chronological age and the predicted age in the HCP test dataset. The left hemisphere model achieved an MAE = 6.70, *r* = 0.93, and *R*^2^ = .87, and the right hemisphere achieved an MAE = 5.54, *r* = 0.94, and *R*^2^ = .89.

**Figure 6.**
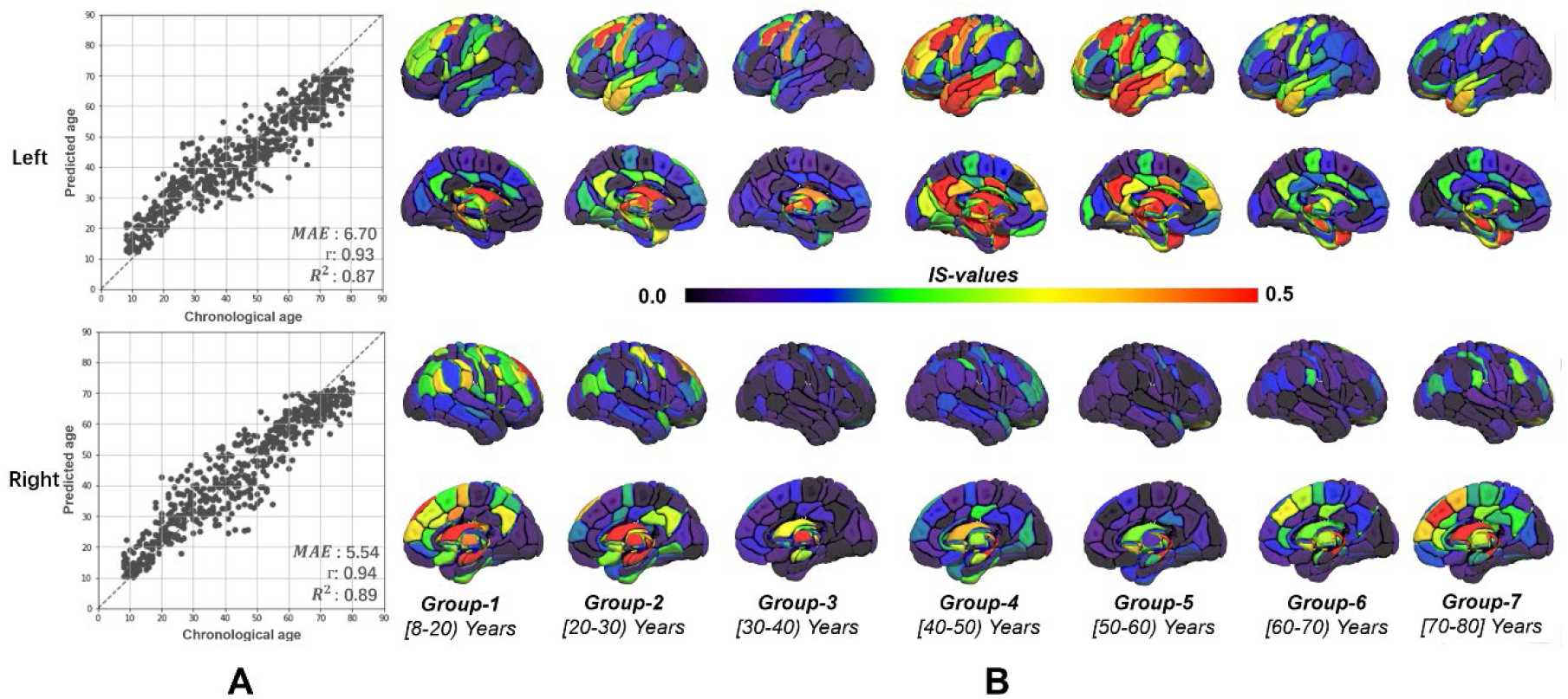
(A) Predictive effects of left and right brain models. (B) Lateralization of the importance of the left suitable brain regions with age. The predictive performance of our model has been verified again in both hemispheres, and the regional lateralization of IS with age was observed. The IS of the temporal lobe area decreased and maintained a stable state in the right hemisphere but increased significantly in the left hemisphere, showing significantly age-related importance in brain aging.

Subsequently, we applied the proposed interpretation method to these two models separately and observed the lateralization of the importance of brain regions with age. We obscured the brain regions with odd IDs in the left hemisphere model while obscuring the brain regions with even IDs in the right hemisphere. Therefore, each model generated the corresponding IS for 123 brain regions. Figure 5(B) shows that the left and right hemispheres had clear lateralization. Similar to the conclusion found for the whole brain, the left and right hemispheres had more important regions during development and aging. However, in the mature stage, we observed distinct lateralization: the essential area decreased and maintained a stable state in the right hemisphere but increased significantly in the left hemisphere (especially in the left temporal lobe area).

## Discussion

In the current study, we trained a lightweight, fully convolutional neural network model for brain age estimation from scratch on a large heterogeneous dataset with uniform and a wide distribution in age and evaluated it on two different datasets (one was unicentric, the other was multicentric, both were also with a uniform distribution in the age ranges). As a result, our best model achieved robustly good performance on both test sets, validating the reliability of the architecture of the model and decreasing the risk of negative effects arising from unstable predictive results in the subsequent analysis.

Then, to further explore the interpretability of the brain age model based on deep learning, we implemented an interpretable scheme combining a network occlusion sensitivity analysis (NOSA) and the human Brainnetome Atlas to interpret the above CNN model, and the results were well verified using different test sets and interpretive methods. We first explored the contribution of brain regions from the whole lifespan by using comprehensive data. We found that the dorsolateral, dorsomedial frontal cortex, anterior cingulate cortex, and thalamus contributed the most to age prediction in the human lifespan. In particular, after occluding these brain regions, the model error changed significantly, reflecting their indispensable position in the brain age prediction process from another perspective. This was also found in the GBP method we implemented and in several recent works [21, 53]. Moreover, considering the definition of brain age as a biomarker of cerebral aging [12], this result simultaneously may reveal a pronounced variation in the corresponding structures from a complete overview of human brain aging, a finding which supports some anatomical studies [54-56] (e.g., our results showed that regions in the thalamus had a dominant influence on brain age prediction, a finding that corresponds to the knowledge that the thalamus is a pivotal brain hub and connects to most parts of the cerebral cortex to form various functional networks [57]). Furthermore, with respect to cognitive changes, the anatomical variability of these regions may be consistent with the continuous process of functional development and decline, including working memory, emotionality, selective attention, and other higher cognitive functions [58-61], findings that also provide evidence for the reliability of our results and interpretative scheme from another perspective. In follow-up experiments, we divided the test set into seven groups to investigate the pattern of variation in the predictive contribution throughout aging and performed the same brain area contribution analysis on each subset. We found that the importance of brain regions changed regularly with age. Brain regions in the frontal lobe became the essential predictors in the developmental and aging stages, whereas the lateral and medial temporal brain regions mainly showed contributions in the aging phase. In the mature stage, most brain regions had no apparent changes, which aligns with what we generally know [62, 63]. In addition, we found that the thalamus had significant contributions at all age stages. These results are meaningful and may imply regional differences in the anatomical or even functional variation with brain aging. Compared to the primary regions in other cerebral structures, the areas in the frontal lobe and the postmedial temporal lobe, which are mainly responsible for higher cognitive functions such as emotionality and memory [64, 65], probably have more age-related changes during the early and late stages of the whole life; these changes could correspond to the early development and the geriatric degeneration of cognition. During the mature stage when the brain is fully developed but may not have begun to age, the age-related changes in anatomy and function are subtle. The continuous variation of the thalamus throughout the human lifespan might be caused by the degree of saliency, which could also be fine-grained but stable and significantly age-related. Moreover, as a hub of most other function networks [66-67], this saliency might be able to explain other age-related changes in the cortex. Based on the analysis above, it is rational to suppose that observing changes in the patterns of brain regions at different ages can help us understand the process of human brain development and aging more deeply. Intriguingly, a similar pattern was not found in the confirmatory experiment in which we used the typical GBP method in the same grouping experiment. This may be because the GBP method is highly related to the model, from which the learned weights dominate the derivatives of each voxel in the model. After the model is trained, the GBP method tends to reveal the invariance learned from training data instead of capturing the variance, which is mainly conditioned on the input [68]. Thus, compared with NOSA, it is no longer very sensitive to subsequent test data, so there is no noticeable change in the results between the groups. However, the NOSA method is sensitive to both the model and the test data, so it can reflect the contributions of different data and capture the changing patterns.

After that, to explore which brain areas would have similar patterns of changes in the brain network contributions throughout the process of brain age prediction, our analysis of the 7-network defined by Yeo showed that the FPN and DMN networks made essential contributions to predicting brain age during the development and aging phases. We also noticed that in the mature stage, the IS values of all the networks were low. Because the model has difficulty learning useful features when there are few changes, the error of the brain age prediction was higher in the mature stage than in other stages. These network-level results could be explained by reasons that are similar to those for the fine-grained assumption described above, and they can be regarded as validation at another level.

In the last experiment, we found lateralization of the importance of brain regions with age. Especially in the mature stage, the left hemisphere has more critical brain regions than the right hemisphere. This insight obtained by our NOSA of brain age model seems to align with some neurobiological studies that showed that the left hemisphere is often dominant in the two brain hemispheres [69-71]. More interestingly, the temporal lobe of the left hemisphere seemed to play an essential role in predicting brain age while the right hemisphere did not. The result could be interpreted as lateralized structural growth and aging in the temporal lobe. Some research has confirmed that the left temporal lobe is different in size from the right and that in most people the left temporal lobe is larger than the right [72, 73], providing further evidence for our interpretation.

Moreover, the result could also be reflected in the age-related variation in functions. Combining this result with the previous function lateralization studies of the temporal lobe [74, 75], it is not difficult to speculate that the perception of verbal and visual information, which is dominated by the left temporal lobe, may be more sensitive to age-related changes and may play an essential role in children’s perceptional development and geriatric functional degeneration. In contrast, the abilities of recognition and non-verbal perception, which are associated with the right temporal lobe, may be modulated by other factors.

Our work demonstrates the feasibility and stability of the NOSA method in interpreting essential areas in brain age prediction and yields an intuitive but efficient paradigm for explaining underlying patterns learned in the predictive model of brain age based on deep learning. This enabled us to verify our model by combining the results with neurobiological findings or even further exploring the actual mechanism of the brain aging process. By exploiting this paradigm, we found the specific brain area contribution change pattern in the process of brain age prediction across the human lifespan. These findings can help us acquire more subtle insights into the changes in cerebral structures and functions. However, the insights need to be further confirmed. Although it can better capture the change process with age than traditional methods, our work still needs improvement. Due to the limited data, there is room for further refinement of our model’s accuracy. We have not yet been able to train a model for each age group to compare whether the results produced are still similar to our findings.

## Conclusion

In conclusion, we implemented a high-precision brain age prediction model on a heterogeneous data set with a large amount of data, uniform distribution, and a wide range of ages. We also used a deep learning interpretability scheme that combined the NOSA method with the human Brainnetome Atlas to explain our model. The proposed method produced similar results on the HCP independent and multi-center tests. It has similarities with the conclusions of other recent studies, which demonstrates the reliability of our results. Our results showed that the dorsolateral, dorsomedial frontal cortex, anterior cingulate cortex, and thalamus made the highest contributions to age prediction during the human lifespan. In subsequent group experiments, we found that different brain regions showed different patterns in brain age prediction at varying age levels and showed significant lateralization. The thalamus plays an especially essential role in the whole aging process. These results reflect the implications of our approach, which could help us better explore the processes of human brain development and aging. In brief, compared with the interpretable methods commonly used in previous work, our approach provides a new interpretable strategy for brain age prediction, which can finely capture the brain regions most associated with brain age prediction and how the contributions of brain regions change with age. In the future, this work can be further refined to improve model performance and can be extended to many other potentially relevant paradigms, such as considering gender differences or applying the model to disease conditions.

## Acknowledgements

This work was partially supported by the Science and Technology Innovation 2030—Brain Science and Brain-Inspired Intelligence Project (Grant No. 2021ZD0200200), the Natural Science Foundation of China (Grant Nos. 82072099 and 62176181), and the Strategic Priority Research Program of the Chinese Academy of Sciences (XDB32030214). The authors appreciate the English language and editing assistance of Rhoda E. and Edmund F. Perozzi, PhDs.

