## Supplementary materials for "Network Occlusion Sensitivity Analysis Identifies Regional Contributions to Brain Age Prediction"

### Supporting Information Text

#### Atlas influence on the NOSA method

Before further exploring the regional importance in the brain, we first examined the influence of the atlas, which is the basis of our network occlusion sensitivity analysis (NOSA) method for defining the cerebral regions. Based on our investigation, we utilized three brain atlases that are all commonly used in cerebral research studies but have differently grained parcellations, specifically, the Anatomical Automatic Labeling (AAL) [1], Brainnetome Atlas (BNA) [2], and Atlas of Intrinsic Connectivity of Homotopic Areas (AICHA) [3]. After that, we used test data and applied the same NOSA method utilizing each atlas separately to quantify the importance of each brain region related to age prediction. The mean result is shown in Figure S1.

The result showed that the importance score (IS) differences between the AAL-defined regions were difficult to identify because almost every region had a high IS value. This may be caused by the low precision and granularity of the AAL parcellation, potentially increasing the anatomical error in the subsequent analysis. However, even though it is a fine-grained atlas with 384 regions, AICHA still did not produce a stable result when using it as the basis for the NOSA method; this was probably due to the AICHA being so fine-grained that it did not yield stable results. In addition, calculating the IS regions in AICHA was time-consuming, resulting in another limitation of the experimental conditions. Finally, only the regions defined by BNA provided a relatively consistent result in every test and thus showed good compatibility with the NOSA method. Moreover, the calculations for the BNA regions were more efficient and had a lower cost than those for AICHA.

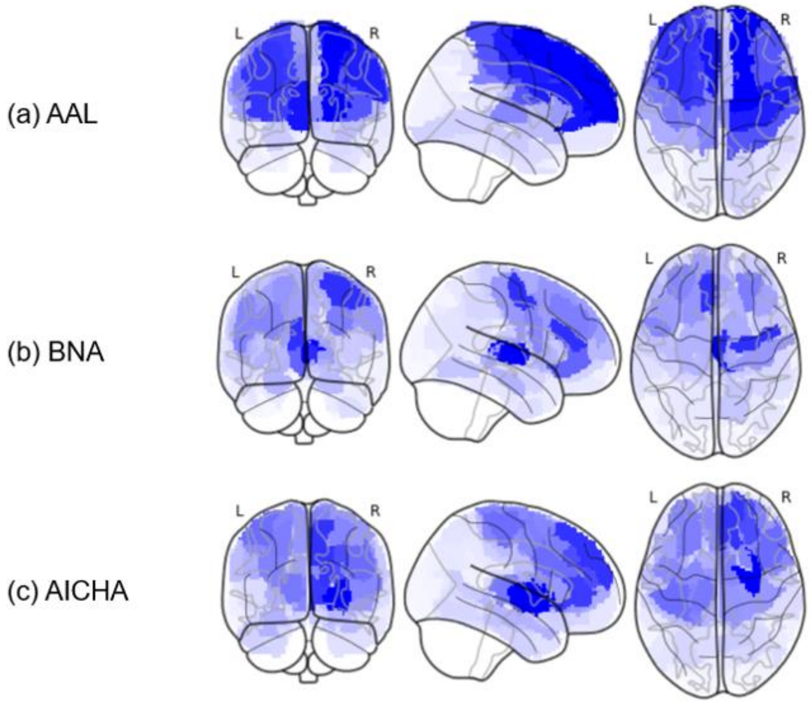


Figure S1. The distribution of IS values on a standard brain template regionally defined by each atlas. The IS differences between the AAL-defined regions were quite small, indicating low usefulness for comparing the changing patterns of the cerebral regions. The result based on the AICHA regions was too fine-grained and lacked stability and the efficiency of the calculation was low. Only the BNA regions showed a pattern of IS variations that was the same in every test and the calculation cost was reasonable.

**Correlation and regression analysis of the changing patterns of age-to-region and region-to-region ISs**

To analyze our findings quantitatively, we performed correlation and regression analyses and aggregated the results by averaging to investigate the age-to-region and region-to-region effects at both the subregional and regionally integrated levels. For the correlation analysis, we employed the Spearman correlation coefficient to evaluate the effect and found that anatomically adjacent regions had similar correlations with age and that the correlation between the regional locations diverged with age. However, compared with the interregional correlation, intriguingly, almost every region showed a similar changing pattern to each of the other regions (compare Fig. S2 and S3), which supported the consistency of our observation in the age groups. We speculate that the different correlative pattern was that the change in regional IS is not dominated by changes in chronological age but depends more on the other regional ISs. In addition, we performed a polynomial regression on the integrated parts and ranked them in order of correlation with age (See Fig. S4). The result showed that the thalamus has both a high average IS value and a strong correlation with age, underscoring its importance in the overall aging process and its changing correlations with other regions.

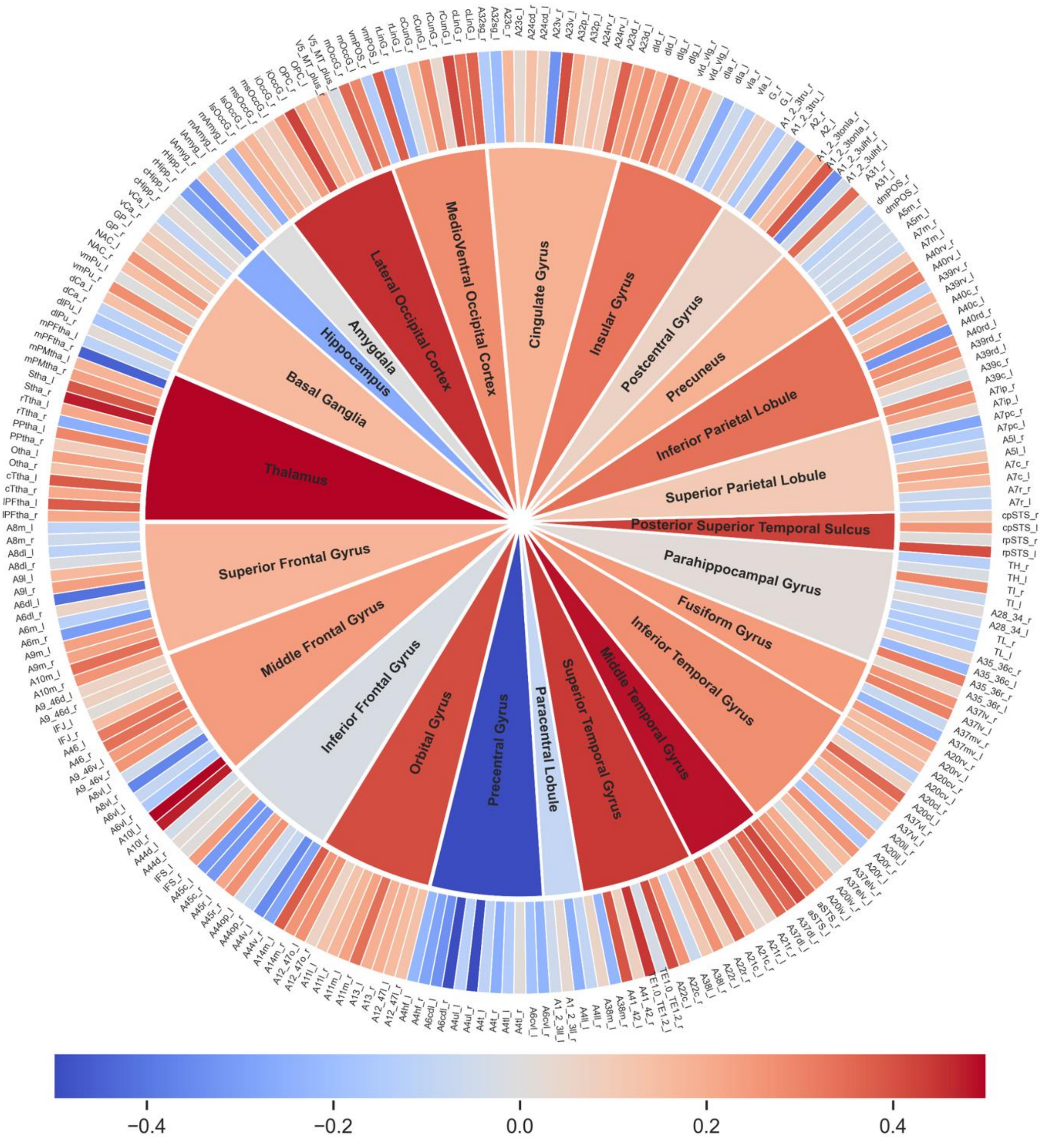


Figure S2. Heat schematic of the correlation between regional IS and age. The outer and inner annuli denote the correlation between subregions and integrated parts, respectively. The red color indicates a positive correlation with age and the blue indicates a negative correlation. We found that the regions in the thalamus had both salient IS values shown in Table 1 of the main text and correlation with age shown here, indicating a high level of consistency.

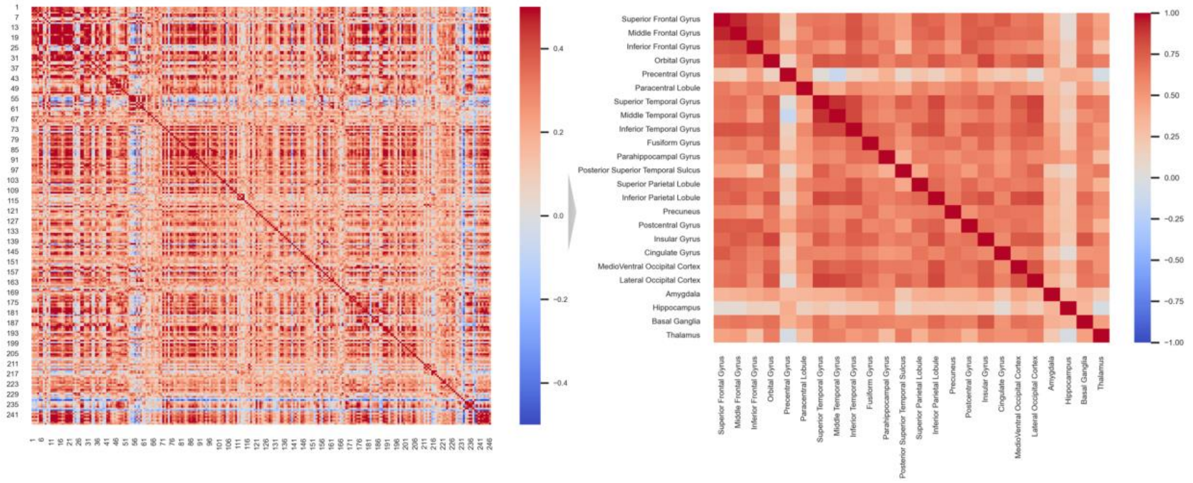


Figure S3. Interregional correlation heatmaps of the correlations. The left map shows the subregional correlation and the right map shows the correlation of the integrated regions. The red and blue colors respectively represent the positive and negative effects. We found that the IS values of the regions generally had a similar change mode, showing a consistent aging pattern throughout the entire life.

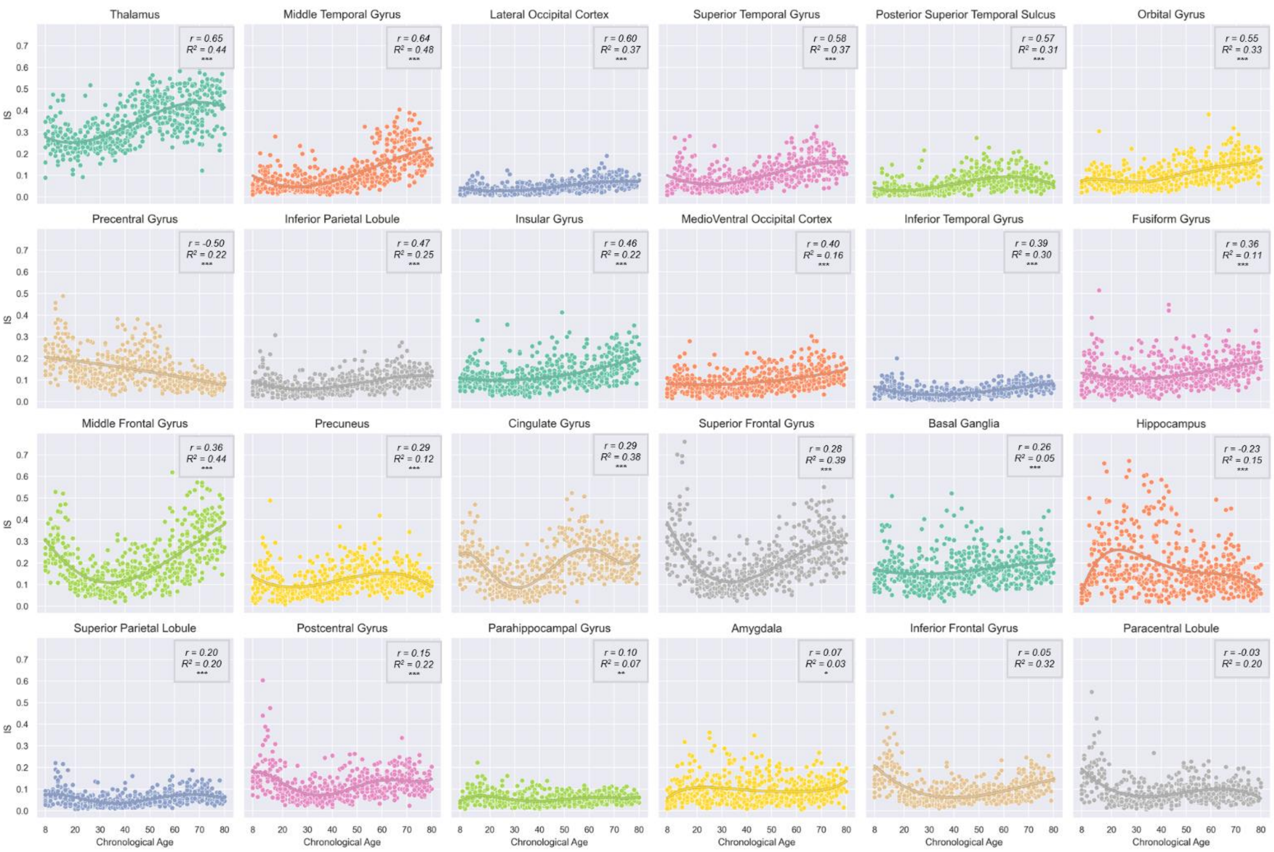


Figure S4. Changing patterns of integrated regions ranked by their correlation with age. The annotations of each subplot are the Spearman correlation coefficient ($r$) and the determination coefficient ($R^2$). All regression functions passed the significance test ($p < .03$). The asterisks below the coefficients show the significance levels of the $r$ ('***': $p < .01$, '**': $p < .03$, '*': $p < .05$). Note that the thalamus had both a high average IS value and a strong correlation with age.
